# BELL: Biomodel Evidence and LLM-based Logic

**DOI:** 10.64898/2026.09.18.721350

**Authors:** Niloofar Arazkhani, Brent Cochran, Natasa Miskov-Zivanov

## Abstract

Building accurate and predictive mechanistic models requires careful biological interaction curation and verification against existing knowledge. When done manually, these tasks become impractical, especially with massive extraction of interactions facilitated by advanced natural language processing methods and large language models (LLMs). We present BELL (Biomodel Evidence and LLM-based Logic), a biocuration support framework that automates evidence retrieval, scoring, and explanation for interaction-level verification. BELL processes each interaction through a five-step pipeline: entity grounding, database ranking, evidence retrieval from seven biological databases, a heuristic four-dimension programmatic scoring, and chain-of-thought explanation with a recommended curator action generated by LLMs. We applied BELL on 210 protein-protein interactions from a curated Glioblastoma Multiforme (GBM) model. Results show that 49.5% of interactions achieved HIGH confidence and 72.4% received a positive curator recommendation, while qualitative flags precisely directed curator attention to evidence gaps. BELL is integrated into the KALIMBA curation platform and available at www.boheme.pitt.edu/Kalimba.

## I. Introduction

Natural language processing (NLP) has expanded the scale at which interactions can be extracted from literature [1, 2], but extraction errors remain common and difficult to detect without manual review [3, 4]. While recent advances in large language models (LLMs) offer an additional layer of reasoning over the extracted knowledge, these models are still prone to hallucinations [5]. At the same time, large scale knowledge extraction is critical for creating comprehensive mechanistic knowledge graphs and executable models of biological systems [6, 7], and the inaccuracies in extraction tools carry real consequences for reasoning about these systems. For example, a single misrepresented or absent interaction can alter predicted network dynamics and produce misleading biological conclusions. Therefore, verification of these networks is critical, as it confirms that individual interactions are supported by biological evidence, a distinct concept from validation, which compares predicted dynamics to experimental outcomes.

Since much of the human interactome is already documented in publicly accessible databases that can be programmatically queried, this offers an opportunity for scalable network verification [8-10]. Systematic assessment of interactions extracted by NLP-based and LLM-assisted tools requires cross-referencing each interaction against multiple databases that differ in coverage, curation standards, and evidence representation, a process that is time-consuming when done manually and difficult to apply consistently at scale. Existing tools such as FLUTE [3] address this by downloading information from databases for offline verification, but this approach requires periodic data updates to remain current, which is not practical as a long-term curation strategy.

To address this gap, we present BELL (Biomodel Evidence and LLM-based Logic), a biocuration support tool that automates the evidence retrieval, scoring, and explanation steps of interaction-level verification. Within BELL, LLMs are prompted using the Chain-of-Thought (CoT) approach. We evaluated the use of BELL on a corpus containing interactions extracted from published literature. Our results confirm that its user-friendly interface substantially reduces the manual effort required for post-extraction curation while providing the evidence transparency needed for informed and reproducible annotation decisions.

## II. Methods

### A. Interaction Processing Pipeline

BELL processes each interaction through a five-step pipeline (Figure 1). The first four steps, namely entity grounding, database ranking, evidence retrieval, and scoring, are fully deterministic ensuring transparency and reproducibility. In the fifth step, Llama 3.1 [11] receives the pre-computed scores as read-only input and generates a natural language CoT explanation along with a recommended curator action (Accept, Accept with enrichment, Flag for review, or Reject). Every step of the evaluation is visible to the curator; the resolved entity names and identifiers, the evidence returned by each database, the computed scores, and the full LLM prompt and response are all displayed in the interface alongside the interaction record.

**Figure 1.**
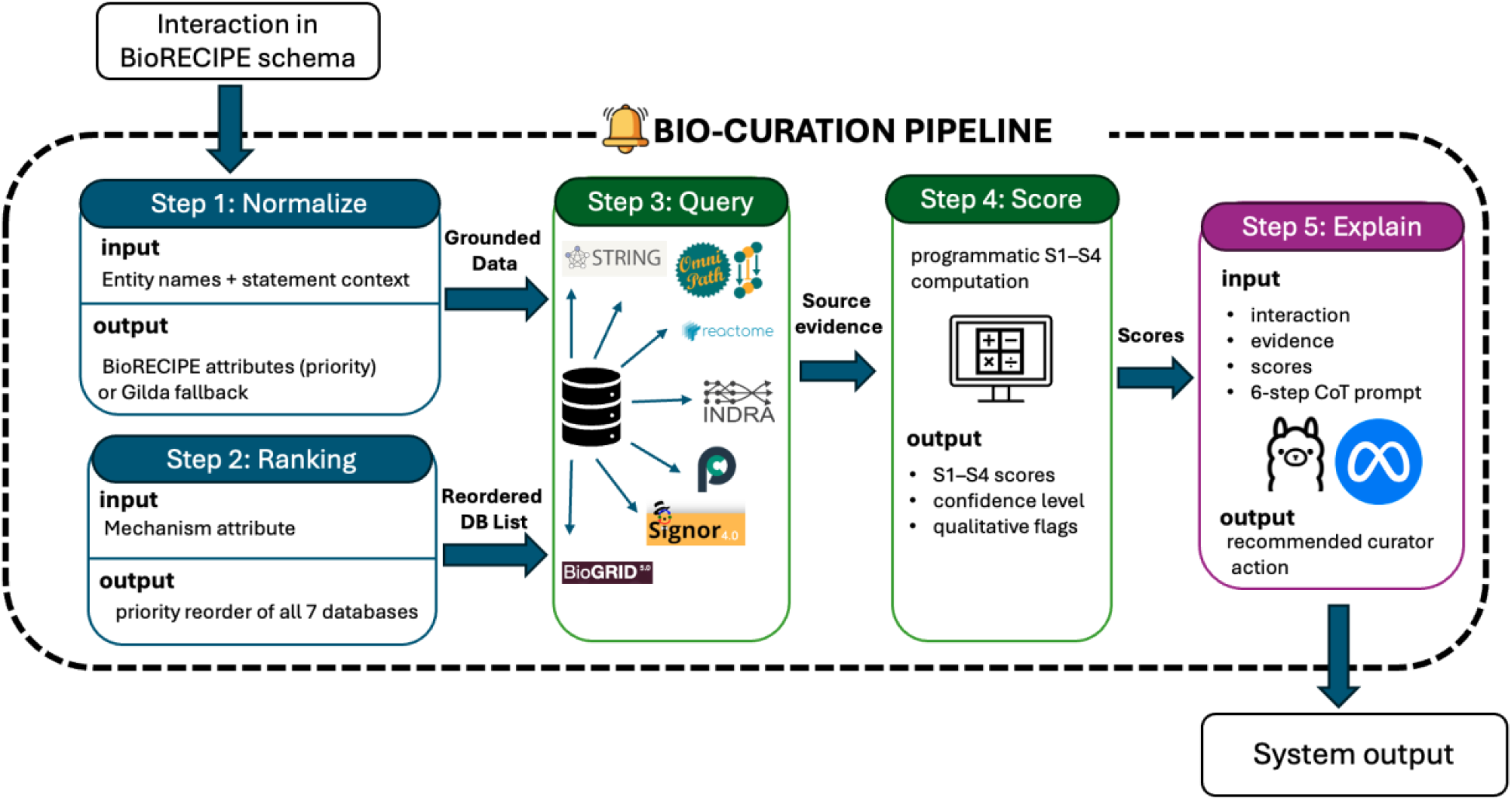
BELL pipeline.

### B. Interaction Representation Format

BELL uses the BioRECIPE representation scheme for the interaction list (BioRECIPE-IL)[12], which documents attributes of the regulator and regulated entities and the interaction between them. For the regulator and regulated entity, attributes include name, type, subtype, unique ID, and compartment. Interaction attributes are Sign, Mechanism, and Connection Type, as well as context attributes such as cell line, cell type, tissue type, and organism, and provenance attributes such as paper ID and source statement from which the interaction is extracted. Interaction direction is included implicitly through assignment of regulator (source) and regulated (target) entities. Detailed documentation for this schema is available in BioRECIPE ReadtheDocs [13]. Other commonly used systems biology formats such as SBML, BioPAX can be converted to and from BioRECIPE using open access converters [14] for broad use of BELL.

### C. Entity grounding

The first step in the pipeline ensures assignment of unique identifiers to entities. When the HGNC Symbol and the Database ID attributes are already populated for both the regulator and regulated entities of an interaction, BELL uses those identifiers directly for querying. For entities without pre-existing identifiers, Gilda [15] resolves the entity name to a canonical identifier using the source statement for disambiguation. The canonical identifier (e.g. HGNC:3236) and HGNC Symbol (e.g. EGFR) are both displayed to the curator in the interface, while entry_name is also used for database queries.

### D. Database selection and evidence retrieval

We selected seven databases to use with BELL: INDRA [16], OmniPath [17], STRING [10], SIGNOR [18], Reactome [8], BioGRID [19], and Pathway Commons [20]. These databases are publicly accessible, widely used in the cell signaling community, and collectively cover the major evidence types relevant to interaction-level verification. Together they span manually curated causal relationships, experimentally detected physical interactions, pathway co-participation, and functional associations, ensuring that evidence is retrieved from complementary sources. Importantly, all seven databases provide free programmatic access via REST APIs, making BELL easily deployable. For every interaction in the input interaction lists, BELL queries all databases uniformly and uses the Mechanism attribute to rank them and reorder the display, prioritizing the databases most likely to contain relevant evidence for that mechanism type. This priority mapping is based on the known specialization of each database: SIGNOR is ranked first for phosphorylation and other post-translational modifications, Reactome for transcriptional regulation, and BioGRID and STRING for physical binding interactions.

### E. Interaction scoring

For each interaction, four scores (*S*_1_-*S*_4_) are computed programmatically from the aggregated evidence. The equations used to calculate them are included in Appendix A: Score definitions.

#### Source coverage

*S*_1_ measures source coverage as a weighted sum of the points from databases where the interaction is found. Weights correspond to database tiers. Tier 1 databases (**INDRA, SIGNOR**) contribute 3 points each as they are based on direct manual curation from primary literature with explicit mechanism annotations. Tier 2 databases (**OmniPath, Reactome, BioGRID**) contribute 2 points each as they provide high-quality but less specific evidence. Tier 3 databases (**STRING, Pathway Commons**) contribute 1 point due to STRING’s reliance on computational inference and Pathway Commons ‘s structural overlap with the other primary databases in the pipeline. To get the final value for *S*_1_, each score is normalized by dividing with the maximum possible weighted sum across queried databases. Overall, higher *S*_1_ indicates broader support from more reliable sources, while lower *S*_1_ shows weak evidence or gaps in database coverage.

#### Attribute match

*S*_2_ measures attribute match across found sources for three interaction attributes: Sign, Mechanism, and Connection Type. For each attribute, a database can report the attribute flags: “1” when the database value matches the declared interaction attribute, and “0” when it contradicts it. If a database’s evidence does not resolve an attribute, or if its data model cannot express that attribute (Table 1), the attribute is marked not applicable and excluded from the average. For each database that found the interaction, the per-database attribute score is the mean of the flags across only the attributes it can report. The final *S*_2_ score is the mean of these per-database scores, considering only databases that found the interaction and reported at least one applicable attribute.

**Table 1.** Attribute coverage by databases.

| Database | Sign | Mechanism | Connection Type<br>(direct/indirect) |
| --- | --- | --- | --- |
| INDRA | ✓ | ✓ | ✓ |
| SIGNOR | ✓ | ✓ | ✓ |
| PathwayCommons | ✓ | ✓ | X |
| OmniPath | ✓ | X | X |
| BioGRID | X | X | X |
| STRING | X | X | X |
| Reactome | X | X | X |

#### INDRA belief score

*S*_3_ is the INDRA aggregated belief score for all retrieved statements supporting the same interaction, reflecting probabilistic confidence derived from literature frequency and source reliability; it is set to 0.0 if INDRA returns no results. *S*_3_ is reported as a supplementary metric but excluded from the confidence threshold (described below), as a score of 0.0 reflects only that INDRA did not return results rather than a lack of supporting evidence overall.

#### Database conflict

*S*_4_ indicates the distribution of databases that provide the supporting and contradicting evidence for the interaction sign, that is, the conflict among databases in interaction sign. In other words, when *S*_4_ = 0, all databases that provide signed interactions agree with the input interaction; when *S*_4_ > 0, there are databases that contradict the sign of the input interaction; when *S*_4_ = 100%, all databases that provide signed interactions contradict the sign of the input interaction.

### F. Qualitative flags

BELL outputs final confidence scores using the following reasoning: HIGH when *S*_1_ ≥ 0.7 and *S*_2_ ≥ 0.5 and *S*_4_ = 0; MEDIUM when *S*_1_ ≥ 0.4 or *S*_2_ ≥ 0.3; and LOW otherwise. To direct curator attention to specific aspects of the evidence, BELL generates qualitative flags. low_source_coverage is raised when *S*_1_ ≥ 0.4, indicating that either none or few databases corroborated the interaction. attribute_match_partial is raised when *S*_2_ ≥ 0.7, indicating that Sign, Mechanism, and Connection Type attributes returned by databases only partially agree with the declared interaction. mechanism_unconfirmed is raised when the input interaction specifies a mechanism but none of the databases that found the interaction returned mechanism-level evidence. sign_conflict_with_input is raised when *S*_4_>0, that is, one or more databases return a sign that contradicts the declared interaction sign.

### G. LLM-based Explanation

While all the previous computations are deterministic and reproducible, in the final step BELL generates CoT explanations using Llama 3.1. The model receives a structured prompt (Appendix B: CoT prompt) containing three sections: the interaction record with source statement, per-database evidence with Connection Type, Sign, and Mechanism annotations, and pre-computed scores *S*_1_-*S*_4_ with interpretive labels. As shown in Figure 2, the model is instructed to reason through six explicit steps including source coverage (*S*_1_), attribute match (*S*_2_), INDRA belief score (*S*_3_), database conflicts (*S*_4_), flags, and recommendation, before producing its output. The model is explicitly constrained from modifying the numeric scores and must conclude with exactly one of four curator actions: ‘Accept’, ‘Accept with enrichment’, ‘Flag for review’, or ‘Reject’. Appendix C: LLM responses shows an example of the LLM’s response suggesting to ‘Accept with enrichment’.

**Figure 2.**
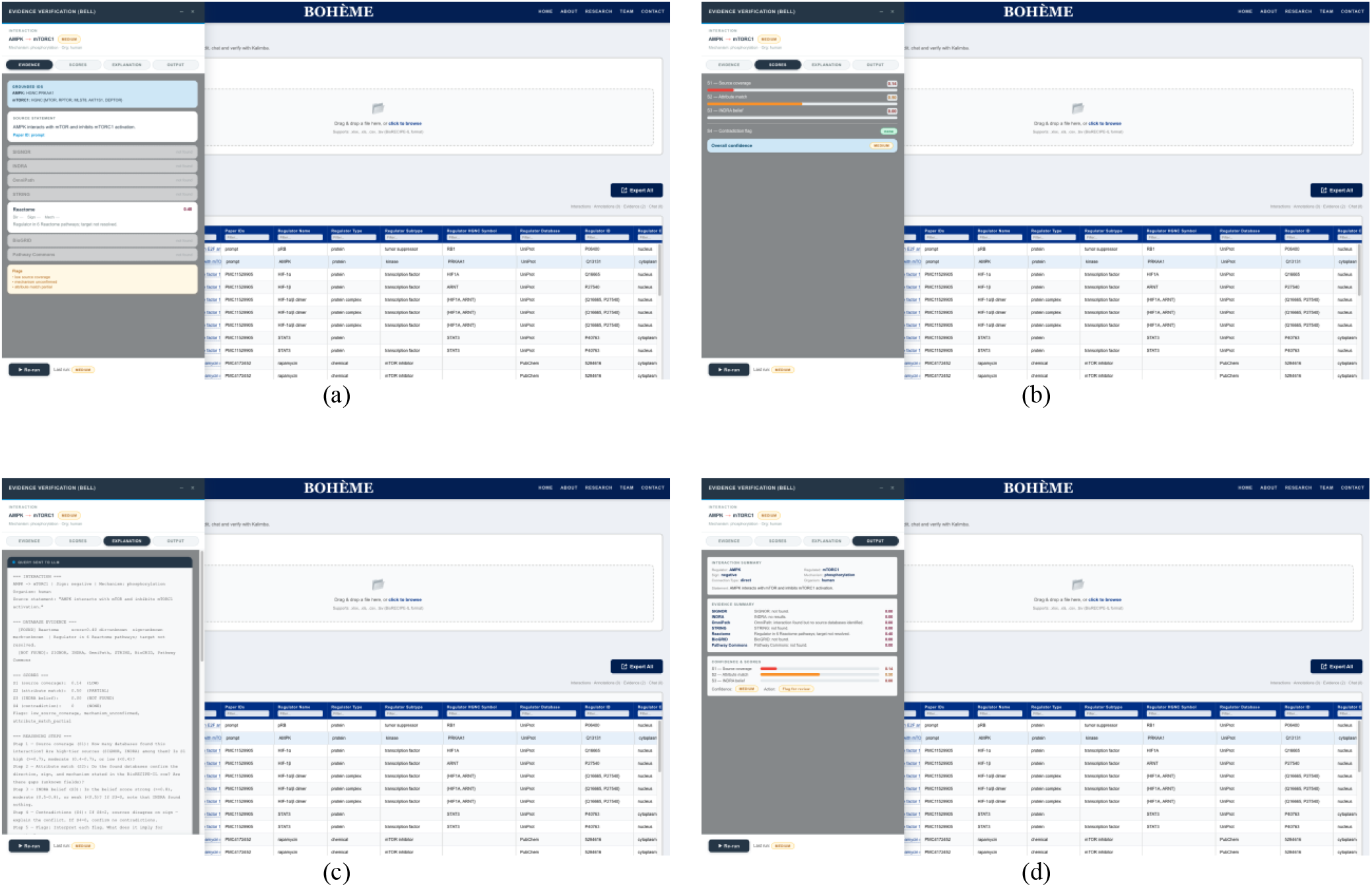
BELL curator interface. (a) Evidence tab displaying per-database query results retrieved from the seven integrated biological databases (b) Scores tab showing S_1_–S_4_ metrics and overall confidence level for a selected interaction. (c) Explanation tab showing the Llama 3.1 chain-of-thought reasoning and recommended curator action. The table supports row-level curator annotation (Accept, Flag, or Reject), inline editing and deletion of individual interactions, and batch export of all evaluated interactions as a downloadable spreadsheet. (d) Output tab summarizing the full interaction record, evidence, scores, and curator status.

## III. Results

In the BELL interface (Figure 2), a curator uploads an interaction list and runs BELL on each interaction. By reviewing the confidence level and recommended action assigned to each interaction, the interactions marked ‘Accept’ or ‘Accept with enrichment’ can be approved quickly, while those with ‘Flag for review’ are brought to attention for closer inspection. The qualitative flags direct the curator to the specific aspect of the evidence that is weak without requiring the curator to read the full explanation for every interaction. For interactions requiring closer inspection, the curator can expand the detail panel to review the full evidence trail and use it to inform their final annotation decision. The curator then annotates each interaction independently using the built-in annotation tool. The fully annotated list can be exported as a spreadsheet for direct use in downstream model construction.

BELL evaluated 210 protein-protein interactions from the curated Glioblastoma Multiforme (GBM) model [21, 22]. The summary of BELL evidence evaluation results (Table 2) shows that the majority received HIGH confidence score (104 interactions, 50%), suggesting strong agreement between the curated interactions and existing knowledge databases. A substantial amount achieved MEDIUM confidence (90 interactions, 43%), indicating partial but not complete database corroboration; however, the GBM model attributes, such as Mechanism, are empty, which biases the results toward MEDIUM even when source coverage is sufficient. Only 16 interactions (8%) received LOW confidence, meaning minimal database support and a need for closer manual inspection using the built-in annotation tool. Moreover, the LLM-based result shows that the largest recommended action for curators is ‘Accept’ (123 interactions, 59%). This means these interactions are well supported by database evidence. A substantial portion of interactions were marked with ‘Flag for review’ (49 interactions, 23%), reflecting MEDIUM confidence scores that need manual curator attention before final acceptance. Twenty-nine interactions (14%) were marked with ‘Accept with enrichment’, representing supported by database evidence but still need additional annotation to fill missing attributes. Only 9 interactions (4%) were recommended as ‘Reject’ due to minimal or absent corroboration in the database. Taken together, 73% of interactions received positive recommendations, while 23% require further curation effort, and just 4% are candidates for removal from the model.

**Table 2.** Summary of BELL evidence evaluation results for the GBM curated interaction set.

| Metric | Category | N | % |
| --- | --- | --- | --- |
| Confidence Score | HIGH | 104 | 50% |
|  | MEDIUM | 90 | 43% |
|  | LOW | 16 | 8% |
| Recommended Action | Accept | 123 | 59% |
|  | Accept with enrichment | 29 | 14% |
|  | Flag for review | 49 | 23% |
|  | Reject | 9 | 4% |
| Quality Flags | Attribute match partial | 59 | 28% |
|  | Low source coverage | 29 | 14% |
|  | Sign conflict with input | 44 | 21% |
|  | Mechanism unconfirmed | 0 | 0% |

Among the three qualitative flags defined in Section II.F, the most repeated one across the evaluated set of interactions is attribute_match_partial, occurring for 59 interactions (28%). This flag reflects cases where the found databases do not fully confirm the Sign, Mechanism, and Connection Type attributes stated in the input row. The absence of Mechanism and Connection Type attributes in the GBM model contributes to the partial score, though a majority of interactions (66%) carried no quality flags at all, indicating sufficient attribute agreement despite the missing attributes. sign_conflict_with_input was raised for 44 interactions (21%), indicating that a meaningful subset of curated interactions has a declared sign that contradicts the sign reported by one or more databases; these interactions need the highest priority for manual re-examination. low_source_coverage was raised for 29 interactions (14%), indicating a subset of interactions found in fewer than 40% of databases.

Figure 3 shows the distribution of *S*_1_, *S*_2_, and *S*_3_ scores across LLM-recommended action categories showed substantial overlap, across the ‘Accept’, ‘Accept with enrichment’, and ‘Flag for review’ categories, most pronounced between ‘Accept with enrichment’ and ‘Flag for review’, suggesting that the LLM recommendation is not fully determined by numeric scores alone but is additionally informed by the qualitative evidence descriptions and reasoning chain provided in the prompt. This behavior is consistent with the intended design of BELL, where the LLM combines numeric scores and database evidence descriptions to produce a recommendation that may capture details not fully reflected in the aggregate scores alone.

**Figure 3.**
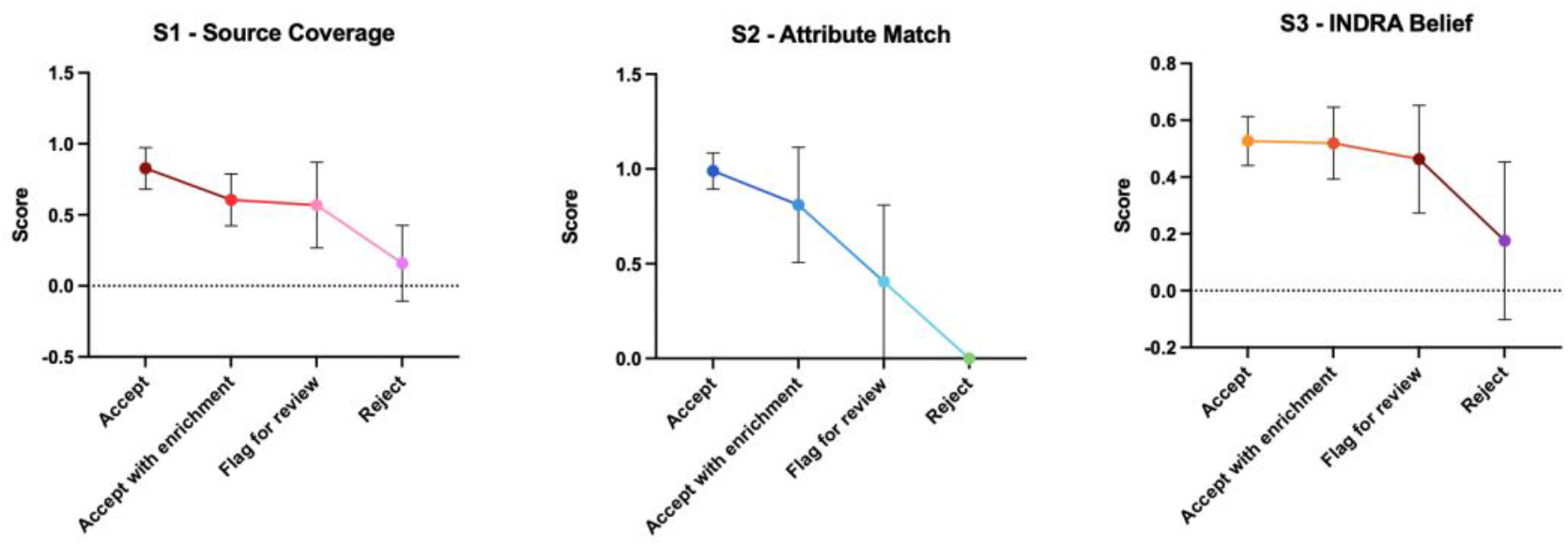
Score distribution by recommended action

## IV. Conclusion

We presented BELL, an automated evidence-scoring pipeline that cross-references biological interactions in BioRECIPE format, against seven major biological databases. The BELL pipeline has two main parts: a deterministic part, which is reproducible and traceable, that generates interpretable confidence scores using a heuristic approach, and a stochastic part, which is based on an LLM with a CoT prompt approach, for recommending an action to the curator based on the results of the deterministic part.

We applied BELL to a 210-interaction GBM dataset; BELL assigned HIGH confidence to 50% of interactions and directed curator attention to a targeted 21% that exhibited Sign conflicts. The attribute_match_partial flag occurred in 28% of interactions and reflected the gap in Mechanism and Connection Type annotations across the current knowledge base. Notably, 66% of interactions showed no qualitative flags, indicating that most curated interactions had had sufficient attribute agreement despite missing annotations. The LLM-generated reasoning layer adds qualitative context beyond numeric scores and provides a curation recommendation.

## Appendix A: Score definitions

For all scores, we define the following:

- Databases/sources: *DB* ∈ {*INDRA, SIGNOR, OmniPath, Reactome, BioGRID, STRING, PCnet*}
- For any interaction *i* and database *DB*, database flag is denoted 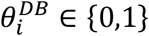 0: database not found as a source of the interaction 1: database found as a source of the interaction

### Score S_1_

Database tiers and corresponding weights *w*:

- Tier I (INDRA, SIGNOR): *w*_I_ = 3
- Tier II (OmniPath, Reactome, BioGRID): *w*_II_ = 2
- Tier III (STRING, Pathway Commons): *w*_III_ = 1 Weighted sum of all sources for interaction *i*:

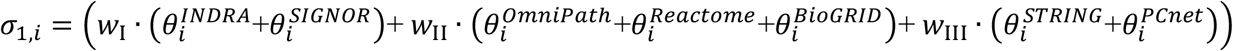

Maximum possible sum when interaction found in all databases:

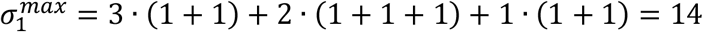

For interaction *i*, score *S*_1_ is: 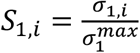

### Score S_2_

We consider the following attributes *A* ∈ {*Sign, Mechanism, Connection Type*}

For interaction *i* and each database *DB* where 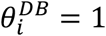, we define the following:

- For each attribute, flag for attribute presence: 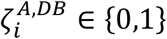 0: value of attribute not reported by database 1: value of attribute reported by database From Table 1, we can determine the value of 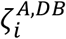 for each attribute and each database.
- Attribute value agreement: 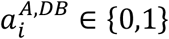 0: contradiction in attribute value (when 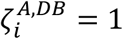) or no attribute value (when 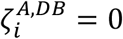) 1: agreement in attribute value (when 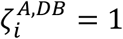)

When there is at least one attribute value reported by database *DB* (i.e., 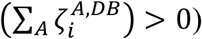), then the normalized database score 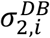 for interaction *i* and database *DB*, across applicable attributes, is calculated as:

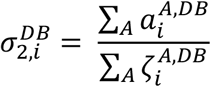

When 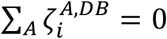, then 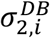 is assigned value 0.

We define *n*_*i*_ as the number of databases where at least one attribute value is found, formally:

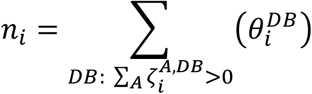

For each interaction *i*, score *S*_2_ is then calculated as an average across all databases that support the interaction and include at least one attribute value:

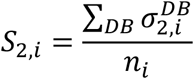

### Score S_4_

For interaction *i, S*_4_ indicates, across all databases where 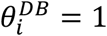 and flag 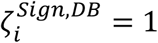, the % of those databases that provide opposite interaction sign. Formally:

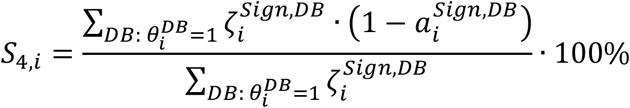

(note that, among seven databases that we use, as shown in Table 1, four provide sign, i.e., 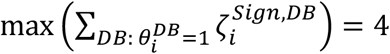

## Appendix B: CoT prompt

**Figure A.**
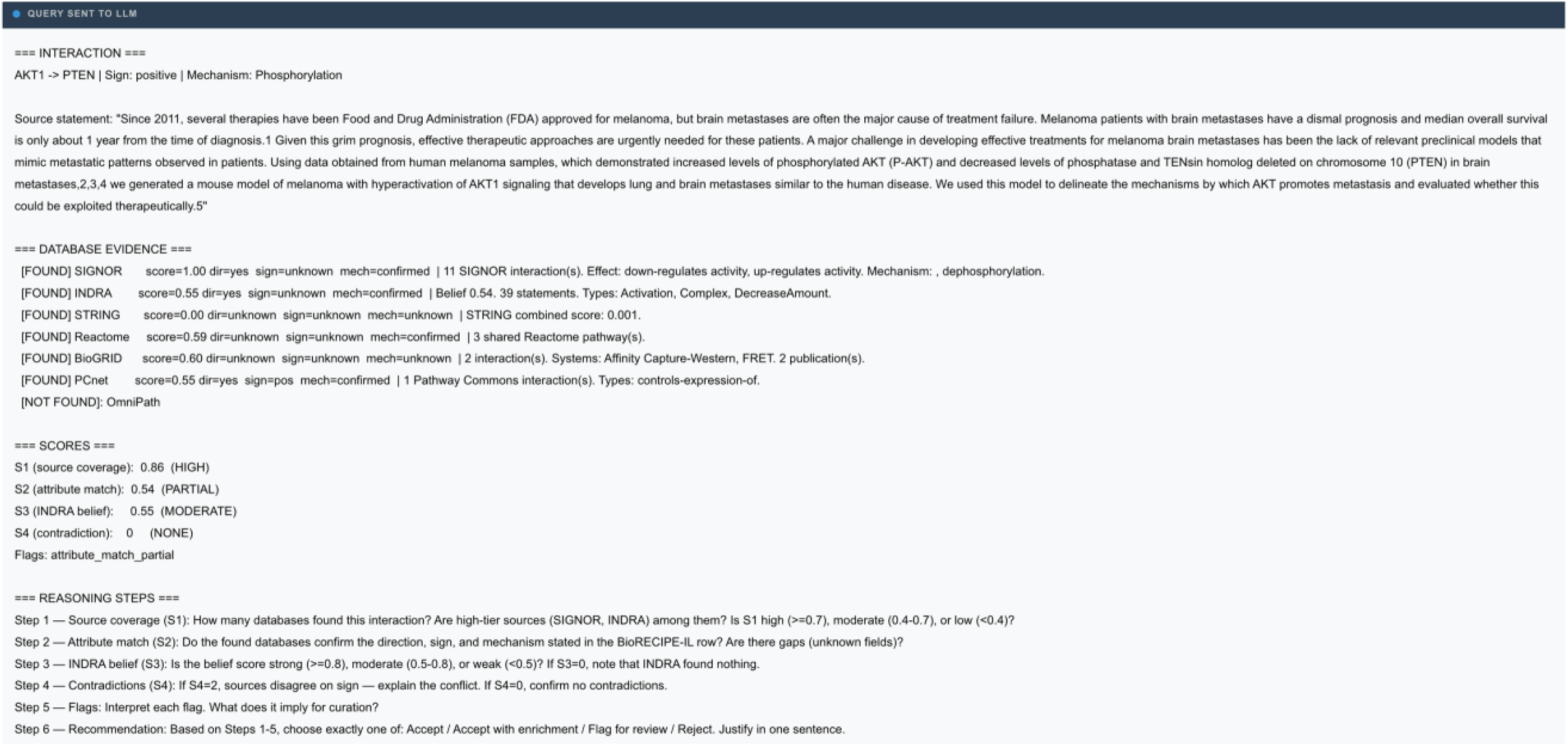
Example chain-of-thought prompt sent to Llama 3.1 for the interaction AKT1→ PTEN (positive, Phosphorylation).

## Appendix C: LLM responses

**Figure B.**
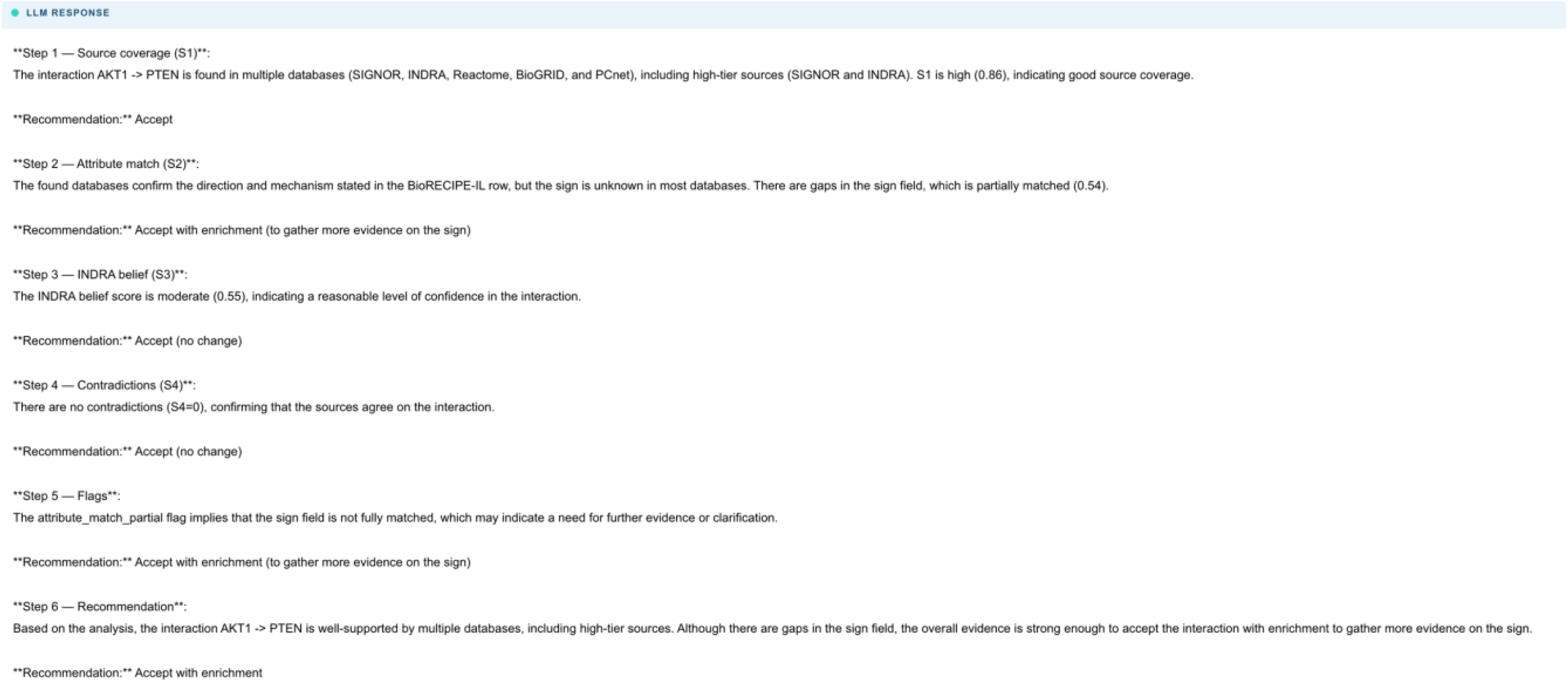
Example of the prompt response in which LLM suggests ‘Accept with enrichment’ for interaction *AKT1*→ *PTEN*.

## Notes

https://boheme.pitt.edu/bell

### Competing Interest Statement

The authors have declared no competing interest.

### Funding

- National Institutes of Health: R01LM014673

